# Breeding for durable resistance in crops: defeated loci may act as Trojan horses compromising the effectiveness of major resistance genes

**DOI:** 10.1101/2025.04.25.650562

**Authors:** Tyrone Possamai, Sabine Wiedemann-Merdinoglu, Marie-Céline Lacombe, Marie-Annick Dorne, Erik Griem, René Fuchs, Jochen Bogs, Raymonde Baltenweck, Komlan Avia, Didier Merdinoglu, Philippe Hugueney

## Abstract

Resistance breeding offers invaluable perspectives for environment-friendly crop protection, but its success may be limited by the breakdown of plant resistance by pathogen strains. This threat is particularly acute for perennial crops, which may be cultivated for several decades. With the increasing use of new varieties carrying multiple major resistance loci, grapevine (*Vitis* spp.) represents a distinctive model to investigate the broad agreement that combining several resistance genes (pyramiding) enhances both resistance efficacy and durability. To this end, grapevine progenies segregating for four resistance loci against *Plasmopara viticola* (*Rpvs*) were used to evaluate the efficiency of single and pyramided major loci when confronted to naive and *Rpv*-breaking pathogen strains. In the context of polygenic resistance, both undefeated and defeated *Rpvs* provided significant quantitative effects. However, interactions between pyramided *Rpvs* were either beneficial, neutral or detrimental to the level of resistance, depending on the loci combination and pathogen strain. In particular, the fact that the presence of defeated resistance loci may compromise the resistance provided by functional major loci has important implications for crops resistance breeding. Thorough phenotypic investigations of pyramiding breeding schemes emerge as a critical step for the effective and durable management of genetic resistances and plant diseases.

## Introduction

Every year, up to 40 percent of world’s agricultural production is lost due to plant pests and diseases [1]. Crop protection is therefore essential for the food security chain, but it largely relies on the recurrent use of agrochemicals, which have adverse human health, environmental and socio-economic impacts [2]. Resistance breeding has been widely used to improve pathogen control while limiting the use of pesticides. However, the breakdown of plant genetic resistance by pathogen strains has become a significant concern for the deployment of plant disease-resistant varieties. The efficient use of genetic resources and resistance durability are particularly important for perennials such as apple (*Malus* spp.) [3], poplar (*Populus* spp.) [4] and grapevine (*Vitis* spp.), which are planted for several decades. Indeed, grapevine has benefited from a worldwide breeding effort to control major diseases such as downy mildew caused by the Oomycete *Plasmopara viticola* [5]. Resistant grapevine varieties have been developed and planted for over 50 years, and, in many countries, cultivated surfaces are currently expanding to reduce the environmental impact of viticulture [5]. This wide array of resistant cultivars, which may harbour multiple pyramided resistance loci, provides a unique model to analyse major resistance genes deployment in crops. Over the past two decades, more than 30 resistance loci to *P. viticola* (*Rpv*) have been identified in *Vitis* spp. [6] (www.vivc.de). For resistance breeding purposes, some major resistance loci have been successfully introgressed in new cultivars and represent very commonly used genetic resources [5,7]. Nevertheless, the high evolutionary potential of *P. viticola* has already led to the development of virulent *P. viticola* strains (*avrRpv-*) that overcome one or more *Rpvs* [8–10]. The construction of polygenic-based resistance has been widely recognised as a strategy to enhance the efficacy and durability of plant resistance [11], and the pyramiding of different *Rpvs* in the same variety is a main objective in grape breeding programs [7]. However, the interactions between pyramided resistance loci are poorly documented, especially when some of these loci have been defeated. Here, we carried out a comprehensive investigation of the efficacy of grapevine major *Rpvs* when confronted with naive and *Rpv*-breaking *P. viticola* strains. Contrasted impacts of defeated major genes in the context of polygenic resistance defines new implications for resistance breeding in grapevine and other crops.

## Results

An offspring of 165 individuals segregating for four resistance loci, *Rpv1, Rpv3*.*1, Rpv10* and *Rpv12*, was assessed in laboratory for the resistance to *P. viticola*, using the standard descriptor OIV452-1 (Fig 1A). The contribution of *Rpvs* to the resistance (Fig 1B) and the resistance levels of the 16 *Rpv* combinations (Fig 1C) were assessed with four different *P. viticola* strain with the following characteristics: one strain was naive, or avirulent, against all the *Rpv* loci [8]; one strain was virulent on *Rpv3*.*1*, or *Rpv3*.*1*-breaking (*avrRpv3*.*1-*) [8]; one strain was *Rpv10*-breaking (*avrRpv10-*); and one was *Rpv3*.*1/Rpv12*-breaking (*avr3*.*1-/12-*) [9].

**Fig 1.**
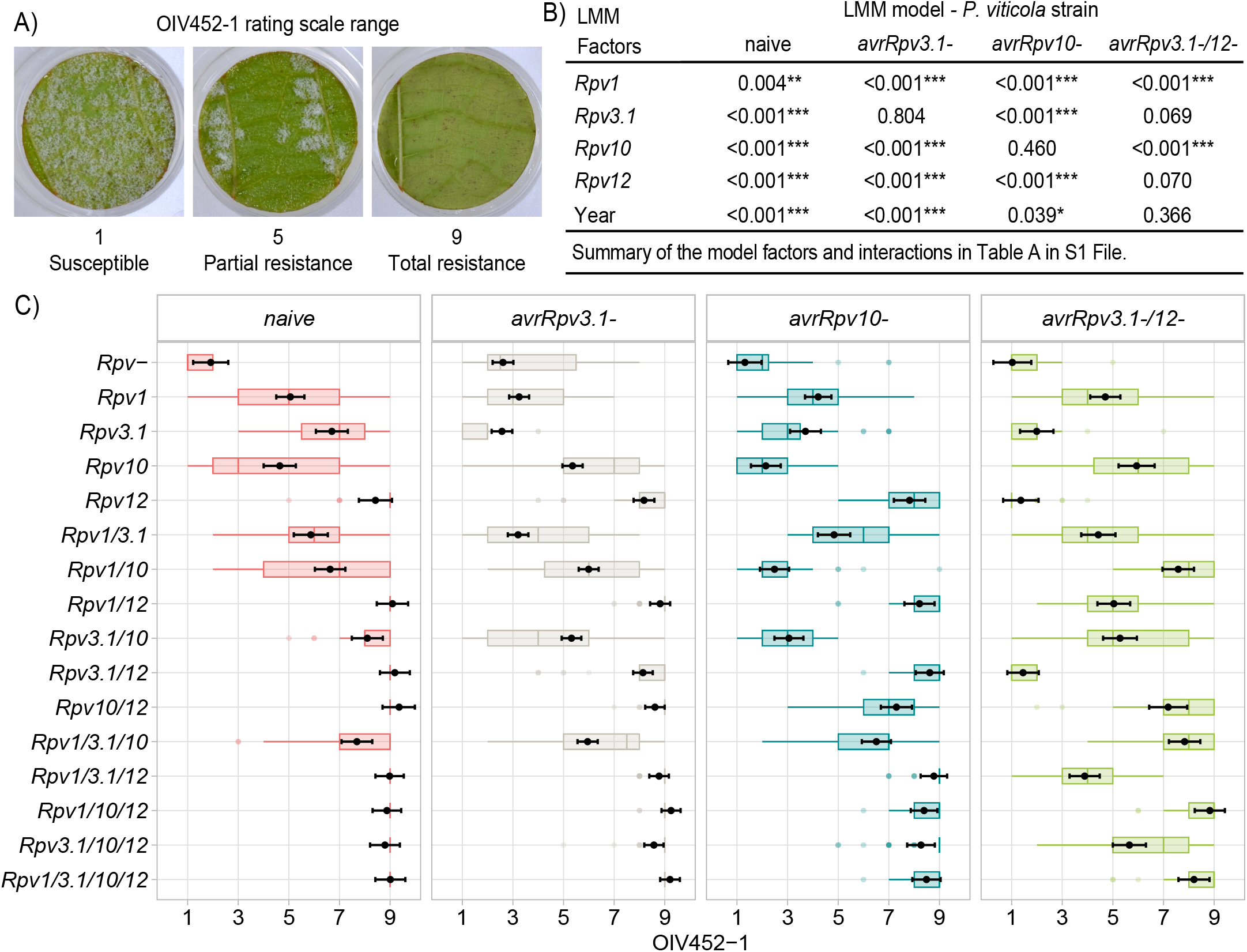
Phenotyping results for the evaluation of the *Rpv*-mediated resistance to *P. viticola*. (A) Grapevine leaf discs illustrating the range of downy mildew symptoms and the associated scoring according to the OIV452-1 descriptor. (B) Table showing significant contribution of grapevine resistance loci to *P. viticola* (*Rpv*) to the resistance based on ANOVA results (p-values: < 0.05 = *, < 0.01 = ** and < 0.001 = ***) performed with the OIV452-1 scores and the optimized linear mixed model (LMM) for the four *P. viticola* strains, naive and virulent (*avr-*) with respect the *Rpvs* tested (see details in Table A in S1 File). (C) Graph showing the distribution of the data (box plots; Table D in S1 File) and the estimate mean with the standard error (black dots and error bars; Table E in S1 File) of the OIV452-1 scores (x-axis) for the studied *Rpv* combinations (y-axis) inoculated with the four *P. viticola* strains (colors). The ANOVA analysis and plots includes all scores from four bioassays and 165 progenies tested (6-12 genotypes/*Rpv* combination) for a total of 36-51 discs/*Rpv* combination/strain (3 discs/strain/genotype/bioassay).

Non-resistant (*Rpv-*) plants exhibited a susceptible phenotype with the naive strain (Fig 1C), associated with a dense pathogen sporulation (OIV452-1 scores in the range of 1-3; Fig 1A). Partial resistances with a less intense sporulation were observed for single *Rpv1, Rpv3*.*1* and *Rpv10* loci (scores between 4 and 7; Fig 1A), and loci-mediated resistance was typically enhanced in pyramided combinations (Table B in S1 File), although the 6 to 12 genotypes representing each *Rpv* combination may exhibit a certain degree of resistance variability (Fig 1C). The *Rpv12* locus, whether used individually or in pyramiding, exhibited the highest and most consistent resistance (Fig 1C), resulting in minimal sporulation of *P. viticola* (scores around 8 and 9; Fig 1A). The three *Rpv*-breaking strains induced susceptible phenotypes in both *Rpv-* plants and those carrying the corresponding defeated *Rpv* genes, in the absence of further pyramiding (Fig 1C). The linear mixed models (LMMs) and ANOVA confirmed that the *Rpvs* significantly contributed to the offspring resistance when they were not overcome (Fig 1B; see details in Table A in S1 File). However, pyramided *Rpvs* combining both effective and defeated genes produced contrasting interactions when challenged with the *Rpv*-breaking strains. Indeed, pairwise comparisons of the *Rpv* combinations showed significant residual resistance benefits of *Rpv10* against the *avrRpv10-* strain and of *Rpv12* against the *avrRpv3*.*1-/12-* strain (Fig 2; Table C in S1 File), observed in one and two pyramided combinations, respectively. Unexpectedly, *Rpv10* also amplified the susceptibility to the *Rpv10*-breaking strain when combined solely with *Rpv1*. Similarly, *Rpv3*.*1* induced detrimental interactions toward the *avrRpv3*.*1-/12-* strain in two *Rpv* combinations, undermining the residual effect observed for *Rpv12* (Fig 2; Table C in S1 File). Nevertheless, the *Rpv3*.*1* locus did not compromise pyramided resistance against the *avrRpv3*.*1-* strain (Table C in S1 File). Altogether, our results highlight unpredictable interactions between defeated and undefeated resistance loci in pyramided combinations.

**Fig 2.**
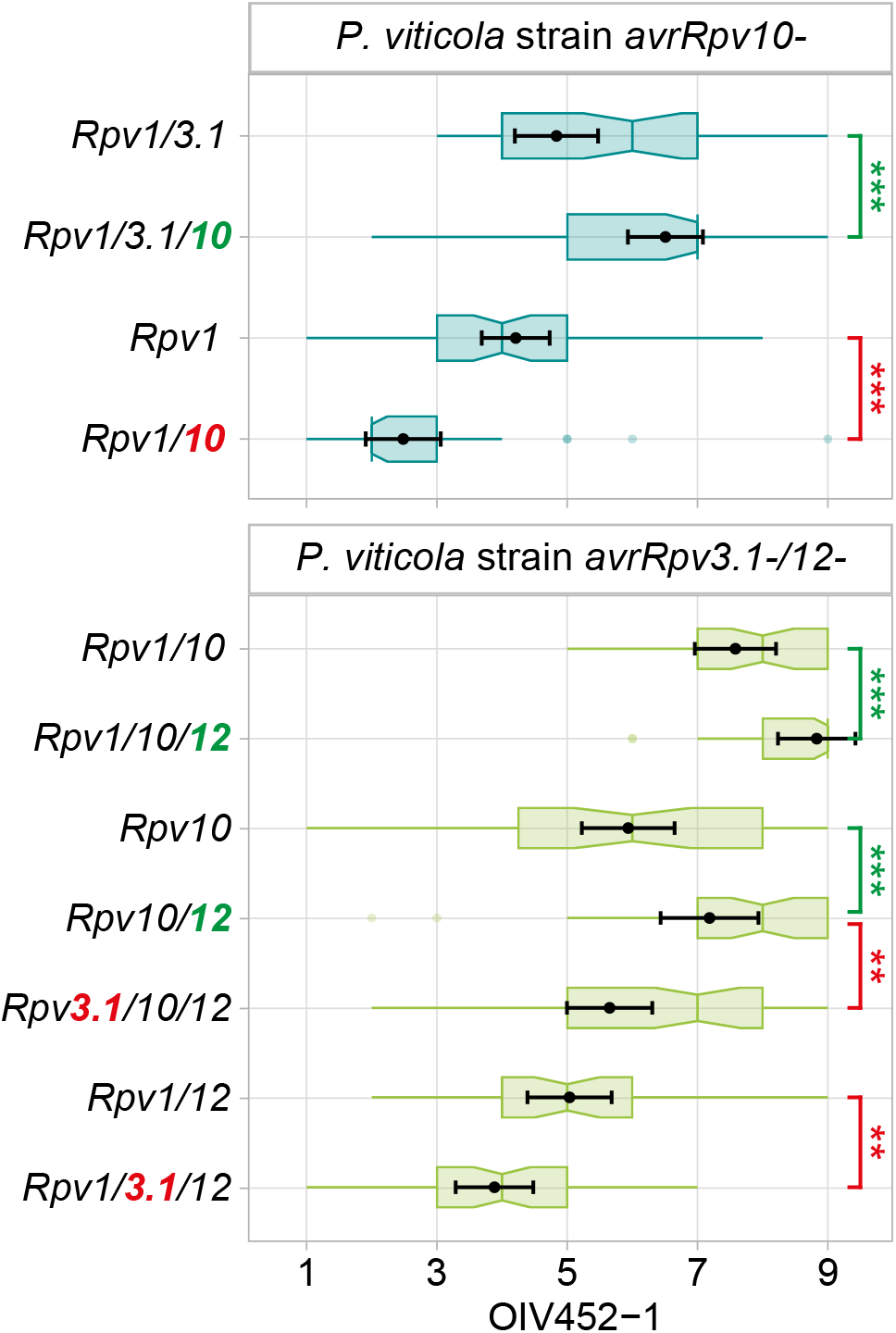
Significant pairwise comparisons between *Rpv* combinations involving defeated loci. Pairwise comparisons of selected *Rpv* combinations displayed in Fig 1C showing significant benefits (green) or downsides (red) of defeated *Rpvs* (coloured) in pyramiding (Tukey’s tests for the OIV452-1 LMM-estimated means and standard errors; p-values with Benjamini-Hochberg correction: < 0.05 = *, < 0.01 = ** and < 0.001 = ***). See data and details in S1 File.

## Discussion

Major loci *Rpv1, Rpv3*.*1, Rpv10* and *Rpv12* have been widely used in grapevine breeding programs [5,7]. The high levels and stability of the resistance to *P. viticola* obtained by combining three or four *Rpv* loci confirmed gene pyramiding as an effective strategy to broaden and enhance plant resistance [11], but infections with *Rpv*-breaking strains also produced unexpected outcomes. The presence of defeated major loci alone or among pyramided loci was most often described as beneficial or neutral for crop resistance [12–15]. However, our results show that in certain genetic combinations, a defeated locus can compromise the resistance provided by undefeated major loci. Although defeated loci have been associated to lower level of quantitative resistance [4], their unpredictable interactions with effective major resistance loci raise important questions about the management of genetic resistance for effective and durable control of fungal diseases. In the context of crop breeding, this suggests that defeated resistance may still be of interest as minor loci to improve and protect effective major loci. However, the potential benefits needs to be confirmed with multiple pathogen strains [4] and through long-term field trials [3,16]. On the other hand, the adverse effects of defeated loci could exacerbate plant susceptibility, thereby compromising the performance and the durability of undefeated major loci. Genetic resistances represent an invaluable natural capital, and it is therefore essential to ensure their sustainable use, even more considering the unpredictable consequences of resistance breakdown. Hence, an integrated approach that combines plant resistance diversity, agronomic practices and agrochemicals remains crucial for optimal pathogen control [17]. For instance, the judicious use of agrochemicals can protect resistant varieties from the emergence of resistance-breaking pathogen strains [18,19], especially in perennials, where pyramiding remains the most viable strategy for the deployment of resistance diversity [17,18,20]. In the context of crop protection, our findings underline that interactions between resistance loci are inherently unpredictable, highlighting the need for comprehensive phenotypic investigations of pyramiding breeding schemes to ensure durable management of genetic resistances and plant diseases.

## Materials and Methods

The biparental grapevine offspring was generated and screened using molecular markers as described in Blasi et al. [21] at INRAE -UMR 1131 SVQV (Colmar, France) in 2023 and 2024. The choice of the parental plants among two advanced breeding lines and the population full sib structure allowed to limit the effect of the genetic background on *Rpv*-mediated resistance. The plants were grown as own rotted plants in 2 l pots in greenhouse, receiving optimal growth conditions (light, temperature and fertilization), regular pruning and the application of sprays for the control of *Erysiphe necator*, the causal agent of grape powdery mildew.

The phenotyping of progenies resistance to *P. viticola* took place under laboratory conditions as described in Wiedemann-Merdinoglu et al. (2022). For the bioassays, the fourth, fifth and sixth leaves from the apex of actively growing shoots were collected and used to produce 2 cm leaf discs. Four *P. viticola* strains were tested with parallel spray inoculations (50,000 sporangia/ml). Leaf discs were visually scored at 6 days-post inoculation (dpi) for the OIV452-1 descriptor [23] as described in Macia et al. [24] but using a nine-classes rating scale (1 = discs completely covered by a dense sporulation – very susceptible to 9 = absence of sporulation – totally resistant). *P. viticola* strains were considered as virulent and the *Rpv* as defeated when the single *Rpv*-strain interaction produced susceptible-like phenotypes as described in Paineau et al. [10].

Four phenotyping bioassays were carried out (two in 2023 and two in 2024) testing 3 discs/strain/genotype/bioassay (most of the genotypes were tested twice) for a final amount of 36-51 discs/*Rpv* combination/strain and 699 discs/strain (Table D in S1 File). A randomisation scheme for the genotypes discs was replicated for each strain and experiment. Two sets of common genotypes (one in 2023 and one in 2024) were used to verify the replicability of the bioassays by t-tests. A linear mixed effect models for each *P. viticola* strain was fitted (‘lmer’ function) and optimised (‘step’ function) by using the ‘lme4’ package [25] in R software [26]. The initial models included the OIV452-1 observations as dependent variable, the *Rpv1, Rpv3*.*1, Rpv10, Rpv12* and the Year as fixed factors with completes cross-interactions, and the experiment-genotype variability as a random factor for the intercept value (Table A in S1 File). Homoscedasticity and normality of the residuals were visually checked in the final models, while ANOVA was used to test the significance of the retained factors and interactions. Estimated marginal means and standard errors of the *Rpv* combinations (Table E in S1 File) were calculated from the optimised models by using the ‘emmeans’ package [27] and used to compare the *Rpv* combinations with Tukey’s tests averaging the data at the year level (p-value with the Benjamini-Hochberg correction; Table B and C in S1 File). All data of the study are included in the article and its supporting information.

## Acknowledgments

We thank Christine Onimus (INRAE UMR1131 SVQV, Colmar, France) for generating the grapevine population; the staff of Unité Expérimentale Agronomique et Viticole (INRAE UE0871, Colmar, France) for maintenance of plant material; and the VEGOIA phenotyping platform (INRAE-Centre Grand Est, Colmar, France), part of the Strasbourg’s University network ‘Cortecs’ (https://cortecs.unistra.fr).

## Funding

This work was supported by the FUNDUR project (ANR-22-CE92-0005; DFG project n° 504993256).

## Author Contributions

PH, JB and DM write the research project and acquired the financial support; PH and SW supervised the research work; TP, SW, MCL, MAD and EG performed the phenotyping bioassays; KA conceived the generation of the offspring; RF isolated the *P. viticola* strains virulent on *Rpv10* (*avrRpv10-*); TP produced the statistical analysis and the visualizations; TP, SW, RB, RF, JB, KA, DM and PH were involved in data and results interpretation; TP and PH drafted the manuscript with input from all authors. The final version of the manuscript has been reviewed, edited and approved by all authors.

## Declaration of interests

The authors declare no competing interests.

## Notes

### Competing Interest Statement

The authors have declared no competing interest.

## References

1. FAO’s Plant Production and Protection Division. FAO; 2022. doi:10.4060/cc2447en

2. European Parliament. Directorate General for Parliamentary Research Services. The future of crop protection in Europe. LU: Publications Office; 2021. Available: https://data.europa.eu/doi/10.2861/086545

3. Lasserre-Zuber P, Caffier V, Stievenard R, Lemarquand A, Le Cam B, Durel C-E. Pyramiding Quantitative Resistance with a Major Resistance Gene in Apple: From Ephemeral to Enduring Effectiveness in Controlling Scab. Plant Disease. 2018;102: 2220–2223. doi:10.1094/PDIS-11-17-1759-RE

4. Dowkiw A, Bastien C. Presence of defeated qualitative resistance genes frequently has major impact on quantitative resistance to Melampsora larici-populina leaf rust in P. × interamericana hybrid poplars. Tree Genetics & Genomes. 2007;3: 261–274. doi:10.1007/s11295-006-0062-0

5. Trapp O, Avia K, Eibach R, Töpfer R. How to deal with the Green Deal – Resistant grapevine varieties to reduce the use of pesticides in the EU. IVES Conference Series. 2023.

6. Possamai T, Wiedemann-Merdinoglu S. Phenotyping for QTL identification: A case study of resistance to Plasmopara viticola and Erysiphe necator in grapevine. Front Plant Sci. 2022;13: 930954. doi:10.3389/fpls.2022.930954

7. Merdinoglu D, Schneider C, Prado E, Wiedemann-Merdinoglu S, Mestre P. Breeding for durable resistance to downy and powdery mildew in grapevine. Oeno One. 2018;52. doi:10.20870/oeno-one.2018.52.3.2116

8. Peressotti E, Wiedemann-Merdinoglu S, Delmotte F, Bellin D, Di Gaspero G, Testolin R, et al. Breakdown of resistance to grapevine downy mildew upon limited deployment of a resistant variety. BMC Plant Biology. 2010. doi:10.1186/1471-2229-10-147

9. Wingerter C, Eisenmann B, Weber P, Dry I, Bogs J. Grapevine Rpv3-, Rpv10- and Rpv12-mediated defense responses against Plasmopara viticola and the impact of their deployment on fungicide use in viticulture. BMC Plant Biol. 2021;21: 470. doi:10.1186/s12870-021-03228-7

10. Paineau M, Mazet ID, Wiedemann-Merdinoglu S, Fabre F, Delmotte F. The characterization of pathotypes in grapevine downy mildew provides insights into the breakdown of Rpv3, Rpv10 and Rpv12 factors in grapevines. Phytopathology®. 2022; PHYTO-11-21-0458-R. doi:10.1094/PHYTO-11-21-0458-R

11. Mundt CC. Pyramiding for Resistance Durability: Theory and Practice. Phytopathology®. 2018;108: 792–802. doi:10.1094/PHYTO-12-17-0426-RVW

12. Pedersen WL, Leath S. Pyramiding Major Genes for Resistance to Maintain Residual Effects. Annu Rev Phytopathol. 1988;26: 369–378. doi:10.1146/annurev.py.26.090188.002101

13. Singh H, Kaur J, Bala R, Srivastava P, Sharma A, Grover G, et al. Residual effect of defeated stripe rust resistance genes/QTLs in bread wheat against prevalent pathotypes of Puccinia striiformis f. sp. tritici. Ishtiaq M, editor. PLoS ONE. 2022;17: e0266482. doi:10.1371/journal.pone.0266482

14. Woo K -S., Newcombe G. Absence of residual effects of a defeated resistance gene in poplar. Forest Pathology. 2003;33: 81–89. doi:10.1046/j.1439-0329.2003.00310.x

15. Taoutaou A, Berindean IV, Chemmam MK, Beninal L, Rida S, Khelifi L, et al. Defeated Stacked Resistance Genes Induce a Delay in Disease Manifestation in the Pathosystem Solanum tuberosum—Phytophthora infestans. Agronomy. 2023;13: 1255. doi:10.3390/agronomy13051255

16. Cowger C, Brown JKM. Durability of Quantitative Resistance in Crops: Greater Than We Know? Annu Rev Phytopathol. 2019;57: 253–277. doi:10.1146/annurev-phyto-082718-100016

17. Zhan J, Thrall PH, Papaïx J, Xie L, Burdon JJ. Playing on a Pathogen’s Weakness: Using Evolution to Guide Sustainable Plant Disease Control Strategies. Annu Rev Phytopathol. 2015;53: 19–43. doi:10.1146/annurev-phyto-080614-120040

18. Carolan K, Helps J, Van Den Berg F, Bain R, Paveley N, Van Den Bosch F. Extending the durability of cultivar resistance by limiting epidemic growth rates. Proc R Soc B. 2017;284: 20170828. doi:10.1098/rspb.2017.0828

19. Taylor NP, Cunniffe NJ. Modelling quantitative fungicide resistance and breakdown of resistant cultivars: Designing integrated disease management strategies for Septoria of winter wheat. Struchiner CJ, editor. PLoS Comput Biol. 2023;19: e1010969. doi:10.1371/journal.pcbi.1010969

20. Zaffaroni M, Papaïx J, Geffersa AG, Rey J-F, Rimbaud L, Fabre F. Combining Single-Gene-Resistant and Pyramided Cultivars of Perennial Crops in Agricultural Landscapes Compromises Pyramiding Benefits in Most Production Situations. Phytopathology®. 2024; PHYTO-02-24-0075-R. doi:10.1094/PHYTO-02-24-0075-R

21. Blasi P, Blanc S, Wiedemann-Merdinoglu S, Prado E, Rühl EH, Mestre P, et al. Construction of a reference linkage map of Vitis amurensis and genetic mapping of Rpv8, a locus conferring resistance to grapevine downy mildew. Theoretical and Applied Genetics. 2011;123: 43–53. doi:10.1007/s00122-011-1565-0

22. Wiedemann-Merdinoglu S, Lacombe MC, Dorne MA, Dumas V, Onimus C, Prado E, et al. Fine monitoring of the effects of grapevine resistance loci on the development of Plasmopara viticola. Caffi T, Rossi V, Fedele G, editors. BIO Web Conf. 2022;50: 02005. doi:10.1051/bioconf/20225002005

23. OIV. Descriptor list for grape varieties and Vitis species, 2nd edn. Office International de la Vigne et du Vin, Paris; 2009. Available: http://www.ov.org

24. Macia FM, Possamai T, Dorne M-A, Lacombe M-C, Duchêne E, Merdinoglu D, et al. Phenotyping grapevine resistance to downy mildew: deep learning as a promising tool to assess sporulation and necrosis. Plant Methods. 2024;20: 90. doi:10.1186/s13007-024-01220-4

25. Bates D, Maechler M, Bolker B, Walker S, Christensen RHB, Singmann H, et al. lme4: Fitting linear mixed-effects models using lme4. Journal of statistical software. 2015;67 (1): 1–48. doi:10.18637/jss.v067.i01

26. R Core Team. R: A language and environment for statistical computing. http://www.R-project.org/. In: R Foundation for Statistical Computing, Vienna, Austria. 2023.

27. Russel L. emmeans: Estimated Marginal Means, aka Least-Squares Means. 2020. Available: https://CRAN.R-project.org/package=emmeans

